# Tagging of water masses with covariance of trace metals and prokaryotic taxa in the Southern Ocean

**DOI:** 10.1101/2023.11.23.568349

**Authors:** Rui Zhang, Stéphane Blain, Corentin Baudet, Hélène Planquette, Frédéric Vivier, Philippe Catala, Olivier Crispi, Audrey Guéneuguès, Barbara Marie, Pavla Debeljak, Ingrid Obernosterer

## Abstract

Marine microbes are strongly interrelated to trace metals in the ocean. How the availability of trace metals selects for prokaryotic taxa and the potential feedbacks of microbial processes on the trace metal distribution in the ocean remains poorly understood. We investigate here the potential reciprocal links between diverse prokaryotic taxa and iron (Fe), manganese (Mn), copper (Cu), Nickel (Ni) as well as apparent oxygen utilization (AOU) across 12 well-defined water masses in the Southern Indian Ocean (SWINGS-South West Indian Ocean GEOTRACES GS02 Section cruise). Applying Partial Least Square Regression (PLSR) analysis we show that the water masses are associated with particular latent vectors that are a combination of the spatial distribution of prokaryotic taxa, trace elements and AOU. This approach provides novel insights on the potential interactions between prokaryotic taxa and trace metals in relation to organic matter remineralization in distinct water masses of the ocean.

## Introduction

The ocean is a dynamical system where hydrological features shape the seascape at multiple scales (Kavanaugh et al. 2014). Hydrographically defined water masses can constrain biogeochemical processes resulting in vertical or horizontal gradients of major nutrients and trace metals (Jenkins et al. 2015). In parallel, the composition of microbial communities that are key mediators in nutrient cycling, varies among ocean basins and along geographical ranges and depth layers (Galand et al. 2010; Agogué et al. 2011; Salazar et al. 2016; Raes et al. 2018; Liu et al. 2019; Sow et al. 2022). Frontal systems, upwelling and mesoscale eddies can structure community composition on a regional scale (Baltar et al. 2010; Lekunberri et al. 2013; Hernando-Morales et al. 2017). Specific hydrographic and biogeochemical properties, among which the concentration of major nutrients, were identified as factors with potential reciprocal influence on these biogeographic patterns in the ocean (Hanson et al. 2012). Trace metals, such as iron (Fe), manganese (Mn), nickel (Ni) and copper (Cu), play crucial roles in microbial growth and metabolism (Morel and Price 2003) and are therefore important micronutrients (Lohan and Tagliabue 2018). In heterotrophic prokaryotes, Fe is essential in the respiratory chain (Andrews et al. 2003), thus Fe availability affects the processing of organic carbon (Fourquez et al. 2014). Mn (II) serves as a cofactor for various enzymes involved in the central carbon metabolism and in antioxidant activity (Hansel 2017). Ni has been identified as an indispensable element for nitrogen fixation (Glass and Dupont 2017) and for chemolithotrophic prokaryotes (Gikas 2008). Cu acts as a cofactor for numerous proteins involved in redox reactions, oxidative respiration, denitrification, and other processes (Argüello et al. 2013). Cu deficiency can affect the growth of some prokaryotic taxa, but certain concentrations of dissolved Cu can also be toxic to prokaryotes or phytoplankton in the ocean (Moffett et al. 1997; Debelius et al. 2011, Posacka et al. 2019).

The biological roles of Fe, Ni and Cu result in nutrient like vertical profiles with low concentrations in surface waters due to biological uptake by auto- and heterotrophic microbes, and increases with depth due to remineralization of sinking material. The magnitude of these uptake and remineralization processes is tightly linked to the composition of the microbial community and its metabolic capabilities. The expected nutrient like profile is not observed for Mn due to the photoproduction of the soluble form of Mn (II) in surface waters and the biologically mediated production of insoluble MnOx at depth (Sunda et al. 1983). Adding to this complexity, transport and mixing largely influence the large-scale distribution of these trace metals (Thi Dieu Vu and Sohrin 2013; Latour et al. 2021; Chen et al. 2023). The GEOTRACES program has made major advances in the determination of the trace metal content of water masses across the global ocean, but the interplay with the microbial community remains to date poorly understood. In this context, the main objective of the present study was to investigate the potential interactive effect between trace elements and microbes, and how these could influence chemical and biological water-mass specific properties across 12 well-defined water masses in the Southern Indian Ocean (SWINGS-South West Indian Ocean Geotraces Section cruise, GEOTRACES GS02 section).

## Materials and methods

### Environmental context

Samples were collected during the SWINGS cruise between January 10 and March 8 2021. The 23 stations sampled for the present study (Fig. 1A) were located in the Subtropical Zone (STZ) (Station 2, 3, 5, 8, 11), Subantarctic Zone (SAZ) (Station 14, 15, 16, 38), the Polar Frontal Zone (PFZ) (Station 21, 25, 31, 33, 36), and the Antarctic Zone (AAZ) (29, 30, 42, 44, 45, 46, 58, 63, 68). Surface water (20m) sampled at each of these stations are assigned to these geographical zones, and the samples below the mixed layer were categorized into 12 water masses according to their physicochemical properties (Fig. 1B).

**Fig 1.**
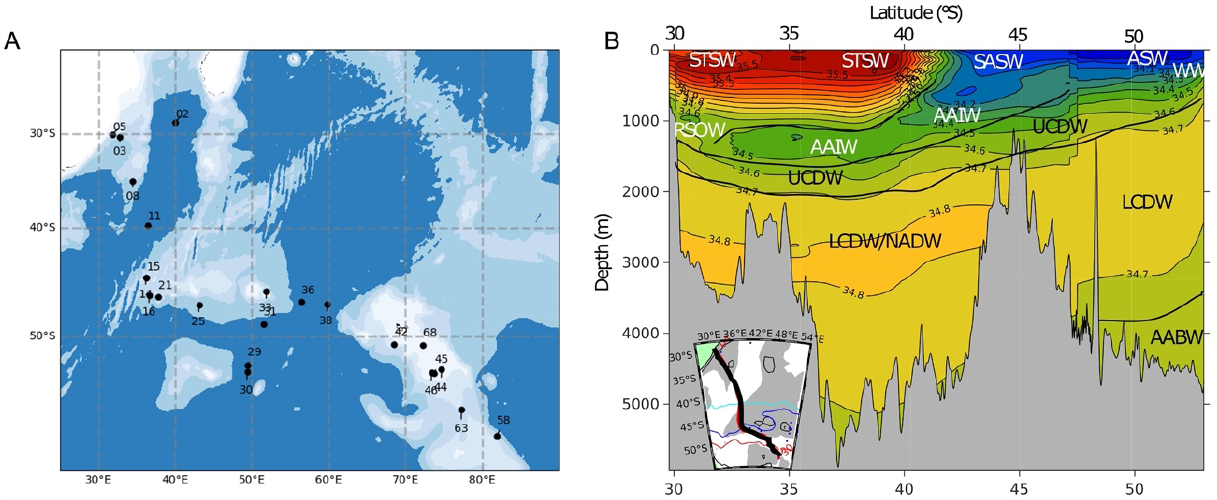
A. Map of stations sampled for the present study during the SWINGS cruise. B. A cross-section (inserted map) showing the vertical distribution of some water masses. STSW, Subtropical Surface Water; SASW, Sub Antarctic Surface Water; ASW, Antarctic Surface Water; WW, Winter water; AAIW, Antarctic Intermediate Water; RSOW, Red Sea Overflow Water; UCDW, Upper Circumpolar Deep Water; LCDW, Lower Circumpolar Deep Water; LCDW/NADW, Lower Circumpolar Deep Water/ North Atlantic Deep Water; AABW, Antarctic Bottom Water. The full list of water masses is provided in Fig. 2.

All seawater samples dedicated to microbial community composition were collected using 12 L Niskin bottles mounted on a rosette equipped with conductivity, temperature and depth (CTD) sensors (SeaBird SBE911plus). Seawater (6L) was sequentially passed through 0.8 μm polycarbonate (PC) filters (47 mm diameter, Nuclepore, Whatman, Sigma Aldrich, St Louis, MO) and 0.22 μm Sterivex filter units (Sterivex, Millipore, EMD, Billerica, MA). The cells concentrated on the 0.8 μm filters were considered particle-attached and those on the 0.22 μm filters as free-living. The filters were stored at -80°C until returned to the home laboratory for DNA extraction. Sample collection, preservation and analyses of prokaryotic abundances, concentrations of dissolved organic carbon (DOC), major nutrients, trace elements and apparent oxygen utilization (AOU) were determined using standard protocols and are described in the Supplemental Methods.

### DNA extraction and sequencing

Total DNA was extracted from the 0.8 μm filters and the 0.22 μm Sterivex filter units using the DNeasy PowerWater Kit (Qiagen) according to the manufacturer’s instructions with a few modifications described in the Supplementary Methods. The V4–V5 region of the 16S rRNA gene was amplified using primer sets 515F-Y (5’-GTGYCAGCMGCCGCGGTAA) and 926-R (5’-CCGYCAATTYMTTTRAGTTT) as described elsewhere (Parada et al. 2016), and PCR amplification was performed as described previously (Liu et al. 2020). 16S rRNA gene amplicons were sequenced with Illumina MiSeq V3 2 × 300 bp chemistry at the platform Biosearch Technologies (Berlin, Germany).

### Data analysis

16S rRNA gene sequences were demultiplexed using the Illumina bcl2fastq v2.20 at the platform Biosearch Technologies (Berlin, Germany). The PCR primers and adapters of 16S rRNA gene sequences were trimmed with cutadapt v1.15. Amplicon sequencing variants (ASV) were produced in R using DADA2 package (v1.24) (Callahan et al. 2016) with the following parameters: truncLen=c(240,210), maxN=0, maxEE=c(3,5), truncQ=2. This pipeline includes the following steps: filter and trim, dereplication, sample inference, merge paired reads and chimera removal. A total of 12,847 unique amplicon sequence variants (ASVs) were obtained from the 172 samples collected (free-living and particle-attached prokaryotes combined). Taxonomic assignment of ASVs were performed using the DADA2-formatted SILVA SSU Ref NR99 138 database (Quast et al. 2012). The number of reads per sample varied between 2,633 and 241,954. Singletons and sequences belonging to Eukaryotes, chloroplasts and mitochondria were removed. To obtain the same number of reads for all samples, the dataset was randomly subsampled to 4,493 reads per sample with the function rarefy_even_depth by the Phyloseq package (v1.40) (McMurdie and Holmes 2013) in R. After subsampling 10,138 ASVs were obtained in total, of which 5,847 ASVs from the free-living fraction (n=76) and 6,461 ASVs from the particle-attached fraction (n=80).

All statistical analyses were performed using the R 4.2.1 version. Non-metric dimensional scaling (NMDS) ordinations were generated based on Bray–Curtis dissimilarity (Legendre and Gallagher 2001) using the ordinate function in the Phyloseq package. Analysis of similarity (ANOSIM) was performed via the vegan package (v2.6) (Dixon 2003) to test for significant differences in microbial communities between water masses. The dendrograms are based on Bray Curtis dissimilarity using the UPGMA algorithm on Hellinger transformed data. To test the association of the free-living prokaryotic community composition and environmental factors, Partial-Least-Squares Regression (PLSR) (Guebel and Torres 2013) analysis with cross-validation was performed using pls v2.8 package (Mevik and Wehrens 2007) in R with the relative abundance of abundant ASVs as the Y variables and the environmental factors as the X variables. Scale-transformation of the data matrix was performed to standardize before data input to the model. The regression coefficients were extract with the function coef by the pls package in R and the heatmap was generated by pheatmap package (v1.0.12) in R. For the identification of indicator ASVs for water masses and surface waters, the IndVal index from the labdsv package (v2.0) (Roberts 2019) in R was used. This index takes into account the specificity, fidelity and relative abundance of the ASVs in the different water masses and surface waters.

## Results and discussion

### Structuring of microbial communities by water masses

In surface waters microbial communities clustered according to geographical zone and frontal system, and in the subsurface the clustering was driven by water mass (Fig. 2). This spatial structuring was significant for both free-living (ANOSIM, R=0.8651, P=0.0001) and particle-attached communities (ANOSIM, R=0.714, P=0.0001). Hierarchical clustering dendrograms and low entanglement values between dendrograms of both size fractions (Fig. S1) further illustrate that the structuring effect of water masses is similar for free-living and particle-attached microbial communities. For a given water mass the composition of the microbial communities was, however, significantly different between size fractions (Fig. S2-4 and Suppl. Results), suggesting that factors that are dependent and independent of size fraction both influence the observed biogeographical patterns. Particles are known to host distinct communities and the nature of the particles can shape the associated prokaryotic assemblages (Baumas and Bizic 2023). Sinking particles were suggested to act as vectors for microbes across the water column (Mestre et al. 2018), an idea that is supported by the about 2-fold lower number of indicator species for a given water mass for the particle-attached (121) as compared to the free-living (213) communities (Fig. S5, Table S1-2, and Suppl. Results). Taken together, these observations point to a complex interplay between processes specific to the particle-sphere and habitat-type independent factors, such as temperature or hydrostatic pressure, to shape the prokaryotic community composition.

**Fig 2.**
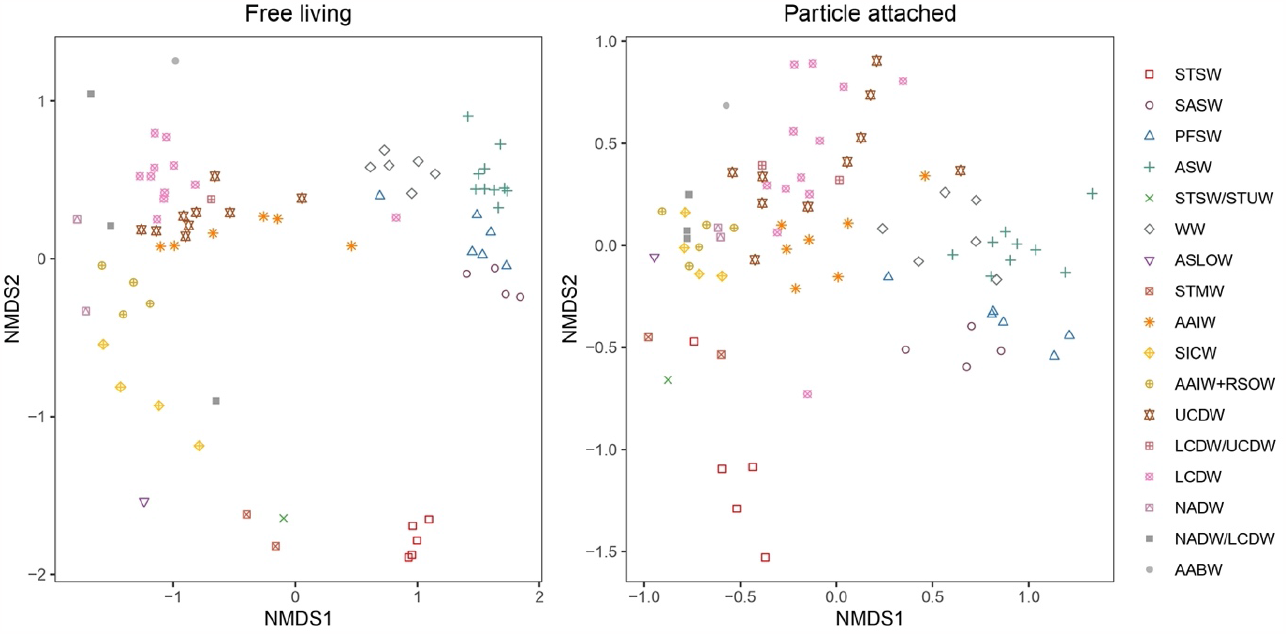
Non-Metric Multidimensional Scaling (NMDS) plots of free-living (FL) and particle-attached (PA) prokaryotic communities based on Bray-Curtis dissimilarity. ANOSIM statistics: FL, R: 0.8651, Significance: 1e-04; PA, R: 0.714, Significance: 1e-04. STSW, Subtropical Surface Water; SASW, Sub Antarctic Surface Water; PFSW, Polar Frontal Surface water; ASW, Antarctic Surface Water; STSW/STUW, Subtropical Surface Water/ Subtropical Underwater; WW, Winter water; ASLOW, Arabian Sea Low Oxygen Water; STMW, Subtropical Mode Water; AAIW, Antarctic Intermediate Water; SICW, South Indian Central Water; AAIW+RSOW, Antarctic Intermediate Water mixed with Red Sea Overflow Water; UCDW, Upper Circumpolar Deep Water; LCDW/UCDW, Lower Circumpolar Deep Water/Upper Circumpolar Deep Water; LCDW, Lower Circumpolar Deep Water; NADW, North Atlantic Deep Water; NADW/LCDW, North Atlantic Deep Water/Lower Circumpolar Deep Water; AABW, Antarctic Bottom Water.

### Microbial ‘biogeo’gradients

Identifying the factors that select for microbial taxa and understanding the potential feedbacks of microbes on the biogeochemical properties of the water mass they thrive in remains challenging. In this context, the role of trace elements in the ocean interior has, to the best of our knowledge, never been considered. To explore the potential reciprocal links between environmental and microbial parameters, we used Partial Least Square Regression (PLSR, or also Projection of Latent Structure Regression). PLSR is a multivariate regression model based on a simultaneous PCA on two matrices which achieves the best relationships between them (Dunn 2020). An advantage of PLSR is that it prevents the bias of co-linearity a facet not taken into consideration by PCA. We carried out PLSR using only those available environmental parameters for which a reciprocal influence can be expected, that are the concentrations of the major nutrients nitrate (NO_3-_) and phosphate (PO_4_^3-^), the trace elements manganese (Mn), iron (Fe), nickel (Ni) and copper (Cu), and Apparent Oxygen Utilization (AOU). To reduce the complexity of the microbial communities, we further considered only abundant prokaryotic taxa, defined as ASVs with a relative abundance of ≥ 5% in at least one sample, resulting in a total of 22 ASVs (Table S3). Because of the stronger clustering of free-living as compared to particle-attached prokaryotes by water masses (Fig. 2) and the availability of the respective dissolved trace metals, we focused our analyses on this size fraction.

The PLSR analysis revealed that the three first latent vectors explained 61%, 16% and 9% of the covariance (Fig. 3A and B; Fig. S6). Therefore, any sample associated with a water mass, which was initially described by 29 variables (7 environmental factors and 22 ASVs) can now be described in a 3-dimensional space. This reduction in complexity facilitates the examination of whether water masses are associated with particular latent vectors which we propose to call microbial ‘biogeo’-gradients (BG). These BGs are a combination of the spatial distribution of environmental factors and ASVs. Our results show that BG1 discriminates deep, cold water masses (UCDW, LCDW, AAIW) (negative signs) from warmer and more saline subtropical waters (STSW/STUW, STMW, ASLOW) (positive signs) (Fig. 3C and D). BG2 mainly discriminates WW (negative sign) (Fig. 3C and E). BG3 provides a partitioning between NADW/LCDW and WW (positive sign) and AAIW and UCDW (negative sign) (Fig. 3C and F).

**Fig 3.**
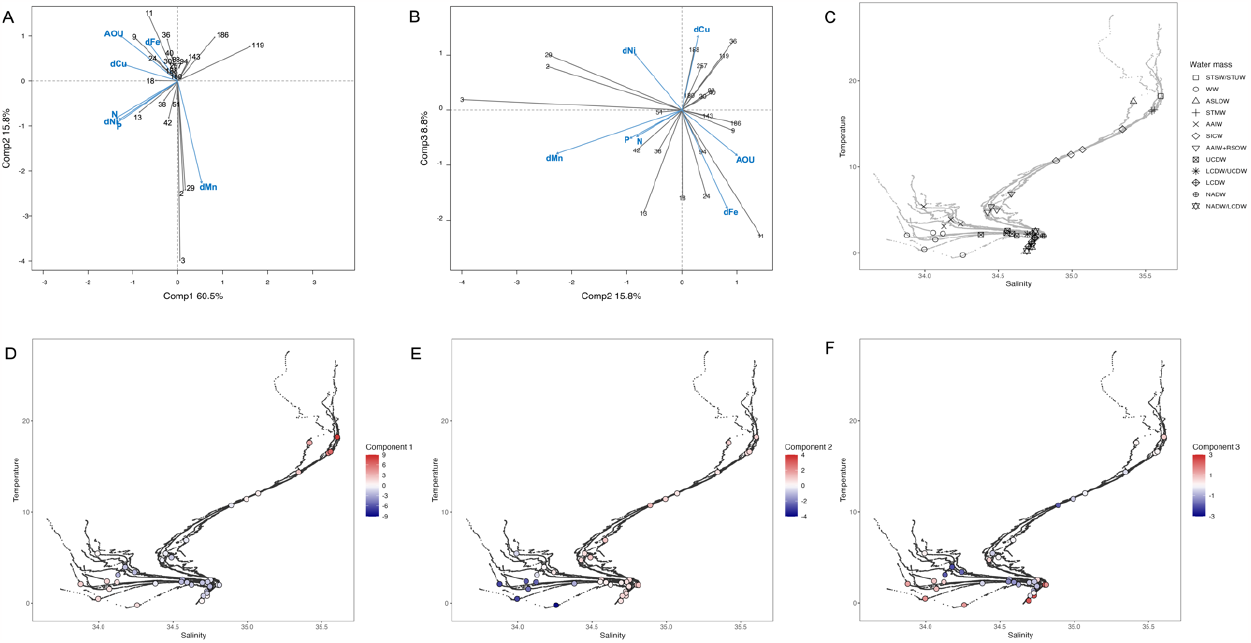
A. Partial least squares regression (PLSR) analysis linking abundant ASVs (relative abundance ≥ 5% in at least one sample) with environmental variables. Blue labels describe the environmental variables (AOU, apparent oxygen utilization; P, phosphate; N, nitrate; dMn, dissolved manganese; dFe, dissolved iron; dNi, dissolved nickel; dCu, dissolved copper) whereas grey labels describe the ASVs (detailed in Fig. 4). Shown are components 1 and 2. B. Components 2 and 3 of the PLSR analysis C. Temperature-salinity diagram and localization of samples collected in different water masses and used for PLSR. D. Temperature-salinity diagram and localization of samples. The color coding corresponds to the first component of scores of samples extracted from PLSR. E. As for panel C, but the color coding corresponds to the second component of scores of samples extracted from PLSR F. As for panel C, but the color coding corresponds to the third component of scores of samples extracted from PLSR.

Physical properties of water masses are set by the conditions at the formation and the subsequent transport and mixing in the ocean interior. These abiotic processes, together with additional biotic transformations contribute to structure on the one hand the distribution of environmental parameters (Fig. S7-9) and on the other hand the distribution of prokaryotic taxa as discussed above (Fig. S3-4 and S10). Our observations that BGs are good descriptors of water masses suggest that they provide clues on the possible interactions between environmental factors and prokaryotic taxa that together contribute to the structuring of latent vectors in the 3-dimensional space.

We discuss in the following these possible reciprocal feedbacks that are the basis of the nature of the BGs. BG1 is dominated by processes linked to remineralization as indicated by the contribution of AOU (Fig. 3A). Therefore, the gradients of the other contributors to BG1 (Fe, Cu, N, P, Ni, and ASVs) across different water masses could be related to this process. Our analysis highlights several ASVs (9, 11, 13, 18, 24) as potential key drivers of remineralization processes (Fig. 3A). BG2 has a more complex structure because it is defined as a gradient with opposite trends between Fe, AOU and the related ASVs (9, 11, 24) and Mn, N, P, Ni and the related ASVs (2, 3, 29). BG3 captures contrasted conditions with opposite gradients between Fe, AOU and the associated ASVs (9, 11, 13, 18, 24, 94), and Ni and Cu associated with another group of ASVs (36, 119, 188, 257) (Fig. 3B and S6). All three BGs are related to remineralization, an observation that is not surprising as this process occurs in all water masses. The regression coefficients, which summarize the information contained in the different BGs, reveal 12 ASVs with a positive relationship with AOU (Fig. 4). The concurrently positive regression coefficients of these ASVs with Fe could indicate that either this element stimulates their metabolic activity and contribution to remineralization or the enhanced supply of Fe by these microbial taxa through remineralization. However, these ASVs could further be partitioned in different groups revealing that Cu is potentially an important discriminating factor. One group of ASVs (11, 13, 18, 24, 38, 94) thrives in low Cu conditions, while another group of ASVs (ASV 9, 30, 40, 88) accommodates with high Cu concentrations. This observation could suggest that the group with negative regressions is either sensitive to the toxicity of Cu or that these ASVs extensively use Cu. Consequently, ASVs belonging to this latter group are potential contributors to the remineralization of Cu.

**Fig 4.**
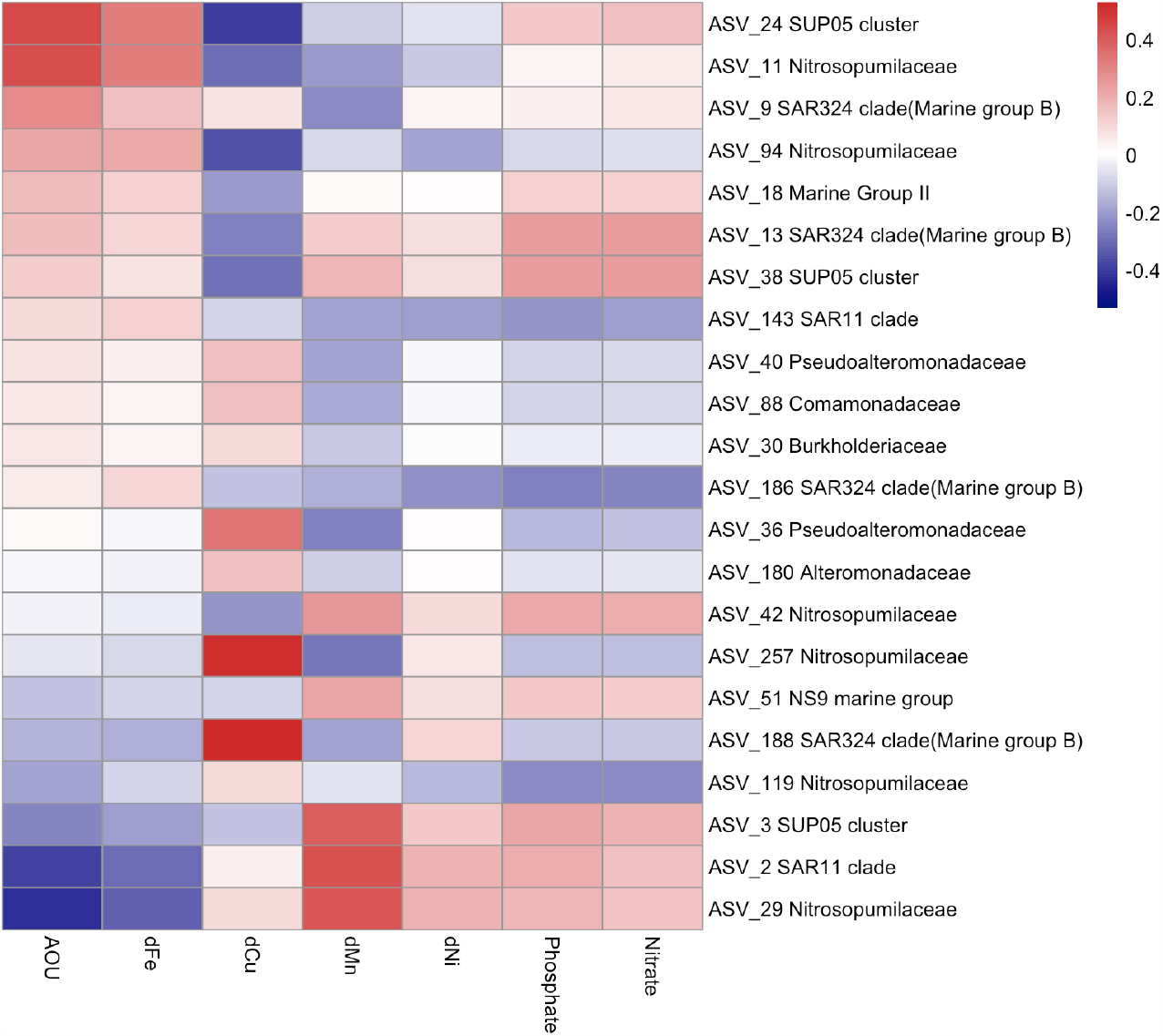
Heatmap based on the regression coefficients of abundant free-living prokaryotes (relative abundance of ASVs ≥ 5% in at least one sample) and environmental variables. The regression coefficients are extracted from the PLSR model.

Negative regression coefficients with AOU were observed with several ASVs suggesting that their activity is decoupled from the remineralization of organic matter. Among these, 3 ASVs (2, 3, 29) had positive regression coefficients with Mn and to a lesser extent with N, P and Ni. These ASVs were highlighted by BG2 that tags WW (Fig. 3E), young water masses with low AOU, typical of HNLC-type waters with high concentrations of N, P and low concentrations of Fe. In the case of Mn, the prokaryotic mediated oxidation of Mn (II) to insoluble Mn (IV) can lead to low Mn concentrations, while photoinduced, organically mediated reduction of Mn (IV, III) can result in high concentrations of this trace element in surface waters (Sunda and Huntsman 1994). This could pinpoint the ASVs with negative regression coefficients (257, 188 and 119) as potential mediators of this reduction (Jones et al. 2020). Another group of ASVs (36, 119, 188, 257) revealed positive regression coefficients with Cu and were significant contributors to BG3, a good marker of NADW/LCDW. The absence of positive regression coefficients with AOU suggests that these ASVs are not Cu remineralizers, but that they are able to thrive in high Cu concentration. This group also contains ASVs that have high negative regression coefficients with Mn.

Our data provide novel insights on the potential interactions between abundant ASVs and trace metals in relation to organic matter remineralization. Among these ASVs, only 7 ASVs were detected by the indicator species analysis (Fig. S5) illustrating the potential of PLSR analysis to identify key microbes if combined with appropriate biogeochemical parameters. Together, these results provide a new view on the parallel distribution of biogeochemical variables and prokaryotic taxa in distinct water masses. Because our results are based on the ASV-level, the limited functional knowledge does not allow to infer the specific pathways involved in trace element cycling by these prokaryotes. However, our results provide the opportunity to identify testable hypotheses on the underlying mechanisms.

We observed that distinct ASVs belonging to the same family revealed opposite regression coefficients with trace elements. This was the case for example of ASVs belonging to *Nitrosopumilaceae*. While ASV 11 and 94 had positive regression coefficients with Fe and negative ones with Cu, ASV 29 and 119 revealed the opposite patterns. *Nitrosopumilaceae* are well-known chemolithoautotrophic ammonia-oxidizers (Qin et al. 2016), but this family also contains members with heterotrophic metabolism (Pester et al. 2011; Aylward and Santoro 2020). Fe- and Cu-availability appears to shape the ecological niches of different strains belonging to this group (Shafiee et al. 2019, 2021). A similar differentiation was observed for ASVs of the SUP05 cluster (ASV 24 and 38 vs ASV 3). Strong positive regressions with Cu were further detected for ASV188 (SAR324 clade, Marine group B), ASV 36 (*Pseudoalteromonadaceae*) and ASV 188 (*Alteromonadacea*). Culture work revealed a range of physiological responses and consequences on cellular carbon metabolism among diverse bacterial strains to Cu gradients (Posacka et al. 2019), illustrating that the requirements of this trace metal or the sensitivities towards its toxicity is highly variable. Insights on the contrasting interplays between trace metals and prokaryotic taxa, including closely related ones, could be gained through the investigation of the gene inventories of the metabolic pathways of interest. Quantifying the respective transporter genes in the water masses where these taxa are abundant and describing the gene repertoire of representative MAGs could be a possible way to further investigate the ecological niches of ASVs in relation to trace metals in future studies.

## Acknowledgements

We thank the captain A. Eyssautier, LDA and GENAVIR officers, engineers, technicians and the crew of the R/V Marion Dufresne for their enthusiasm and their professional assistance during the SWINGS cruise. The SWINGS project was supported by the Flotte Océanographique Française (10.17600/18001925), Agence Nationale de la Recherche (ANR 19-CE01-0012), CNRS/ INSU (Centre National de la Recherche Scientifique/Institut National des Sciences de l’Univers) through its LEFE actions, Université de Bretagne Occidentale, and IsBlue project, Interdisciplinary graduate school for the blue planet (ANR 17-EURE-0015) and co-funded by a grant from the French government under the program ‘Investissements d’Avenir’ embedded in France 2030. We thank Carina Bunse who shared the code for PLSr. We thank the French Institute of Bioinformatics (IFB; https://www.france-bioinformatique.fr) for providing computing resources. This work is part of the PhD thesis of R.Z. supported by the China Scholarship Council (CSC; No. 202006220057).

## Declaration of Competing Interest

The authors declare no competing interests.

## Appendix

A. Supplementary data

Supplementary data to this article can be found in the online version of this article.

## Notes

http://www.obs-vlfr.fr/proof/php/SWINGS/swings_PRECRUISE.php.

### Competing Interest Statement

The authors have declared no competing interest.

